# Mitochondrial mutations in *Caenorhabditis elegans* show signatures of oxidative damage and an AT-bias

**DOI:** 10.1101/2021.05.03.442463

**Authors:** Gus Waneka, Joshua M. Svendsen, Justin C. Havird, Daniel B. Sloan

## Abstract

Rapid mutation rates are typical of mitochondrial genomes (mtDNAs) in animals, but it is not clear why. The difficulty of obtaining measurements of mtDNA mutation that are not biased by natural selection has stymied efforts to distinguish between competing hypotheses about the causes of high mtDNA mutation rates. Several studies which have measured mtDNA mutations in nematodes have yielded small datasets with conflicting conclusions about the relative abundance of different substitution classes (i.e. the mutation spectrum). We therefore leveraged Duplex Sequencing, a high-fidelity DNA sequencing technique, to characterize *de novo* mtDNA mutations in *Caenorhabditis elegans.* This approach detected nearly an order of magnitude more mtDNA mutations than documented in any previous nematode mutation study. Despite an existing extreme AT bias in the *C. elegans* mtDNA (75.6% AT), we found that a significant majority of mutations increase genomic AT content. Compared to some prior studies in nematodes and other animals, the mutation spectrum reported here contains an abundance of CG→AT transversions, supporting the hypothesis that oxidative damage may be a driver of mtDNA mutations in nematodes. Further, we found an excess of G→T and C→T changes on the coding DNA strand relative to the template strand, consistent with increased exposure to oxidative damage. Analysis of the distribution of mutations across the mtDNA revealed significant variation among protein-coding genes and as well as among neighboring nucleotides. This high-resolution view of mitochondrial mutations in *C. elegans* highlights the value of this system for understanding relationships among oxidative damage, replication error, and mtDNA mutation.

## INTRODUCTION

Mitochondrial genomes (mtDNAs) of most animals have mutation rates about one order of magnitude greater than their corresponding nuclear genomes (Wolfe *et al*. 1987; Denver *et al*. 2000, 2004; Havird and Sloan 2016; Allio *et al*. 2017). While rapid mtDNA mutation rates of metazoans have proven useful for understanding the divergence of closely related species (Bernt *et al*. 2013; Yang *et al*. 2016), they also pose a serious challenge for organismal fitness (Gemmell *et al*. 2004). In humans, mtDNA mutations cause genetic disorders (Longley *et al*. 2005; Marni J. and Soondhiemer 2010), are associated with numerous cancers (Gorelick *et al*. 2021), and accumulate with age (Kennedy *et al*. 2013). Further, individuals suffering from age-related diseases such as Parkinson’s and Alzheimer’s display increased mtDNA mutations compared to healthy individuals (Monzio Compagnoni *et al*. 2020). The mechanisms underlying high mtDNA mutation rates in metazoans remain the subject of ongoing research and debate.

Historically, elevated mtDNA mutation rates have been hypothesized to be driven by oxidative damage (Harman 1972; Miquel *et al*. 1980; Richter *et al*. 1988; Shigenaga *et al*. 1994) from reactive oxygen species (ROS), which are abundant biproducts of electron transport in the mitochondria (Murphy 2009). Several studies measuring mtDNA mutations in metazoans have raised doubts about the oxidative damage hypothesis. Specifically, genetic backgrounds with increased ROS do not show a detectable increase in mtDNA mutations (Itsara *et al*. 2014), nor do those with deficiencies in oxidative damage repair machinery (Halsne *et al*. 2012; Itsara *et al*. 2014; Kauppila *et al*. 2018). In addition, the mtDNA mutation spectra from humans is relatively deplete of CG→AT transversions (Kennedy *et al*. 2013), a substitution class considered a hallmark of oxidative damage (Cheng *et al*. 1992; Kirkwood and Kowald 2012).

Alternatively, high metazoan mtDNA mutation rates may be driven by replication errors or deficiencies in mtDNA repair machinery (Longley *et al*. 2005; Szczepanowska and Trifunovic 2015; DeBalsi *et al*. 2017; Hood *et al*. 2019). The replication error hypothesis is supported by *in vitro* mtDNA replication assays with Pol γ (the metazoan mtDNA polymerase), which recapitulate most (83%) of the mutational hotspots detected in a portion of the human mitochondrial genome through phylogenetic methods (Zheng *et al*. 2006). The *in vivo* mtDNA mutational spectra of humans, fruit flies, and mice, are all dominated by CG→TA transitions, which are commonly attributed to Pol γ error (Zheng *et al*. 2006; Kennedy *et al*. 2013; Itsara *et al*. 2014; Melvin and Ballard 2017; Arbeithuber *et al*. 2020). However, because cytosine deamination (to uracil) is likely a major cause of CG→TA transitions in metazoan mtDNAs, the high abundance of these substitutions may reflect a complex relationship between single-stranded DNA damage, failed DNA repair, and replication error. Deamination of cytosine can occur via spontaneous hydrolysis (Nabel *et al*. 2012), but oxidative damage can also play a role by creating modified bases that are more prone to deamination and/or less accessible to repair pathways (Kreutzer and Essigmann 1998). It is not clear whether cytosine deamination in metazoan mtDNAs is driven by spontaneous hydrolysis, oxidative damage, or a combination of the two. If deaminated cytosines are not repaired through base excision repair (BER), Pol γ frequently incorporates adenine opposite of uracil during DNA replication (Zheng *et al*. 2006). In a successive round of DNA replication, the pairing of thymine with adenine completes the double-stranded CG TA transition (Nabel *et al*. 2012). The role of deamination in CG TA transitions in metazoan mtDNAs is supported by strand asymmetry analyses, which have revealed significantly higher frequencies of C→T than G→A changes on mtDNA strands that spend increased time in single-stranded states during mtDNA replication (Kennedy *et al*. 2013; Itsara *et al*. 2014; Ju *et al*. 2014; Arbeithuber *et al*. 2020). Such asymmetric single-stranded exposure likely explains why the C→T change occurs on the “minor strand” in 91% of CG→TA transitions in fruit fly mtDNA (Itsara *et al*. 2014). Interactions between damage and polymerase fidelity can be difficult to untangle. For example, a recent study showed the *in vitro* proofreading activity of human Pol γ decreased when oxidative stress was applied to the enzyme (Anderson *et al*. 2020).

A major obstacle in understanding causes of metazoan mtDNA mutations stems from the difficulty of detecting rare mutation events and removing the biasing effects of selection. Mutation accumulation (MA) experiments (Katju and Bergthorsson 2019) have been used to address this challenge in fruit flies (Haag-Liautard *et al*. 2008; Keightley *et al*. 2009), mice (Uchimura *et al*. 2015), water fleas (Xu *et al*. 2012) and most extensively, nematodes (Denver *et al*. 2000; Howe *et al*. 2010; Molnar *et al*. 2011; Konrad *et al*. 2017; Wagner *et al*. 2020). MA studies remove the filtering effects of natural selection by bottlenecking experimental lines through randomly selected individuals for successive generations. Under these conditions, non-lethal mutations are expected to accumulate and can be assayed through genome resequencing of the final MA generation. However, while MA experiments have proven extremely useful for studying mutation (Lynch *et al*. 2016), this approach poses special challenges for measuring mtDNA mutations (Schaack *et al*. 2020). Mitochondrial genomes remain multicopy throughout the entirety of metazoan germline development (Bratic *et al*. 2010; Wai *et al*. 2010), giving selection the opportunity to act on competing mtDNAs within an individual (Fan *et al*. 2008), even during MA experiments (Schaack *et al*. 2020). The polyploid nature of mtDNAs means that new mutations are born into a low frequency, heteroplasmic state (multiple haplotypes within the same mitochondria or individual). New mtDNA mutations must therefore rise in frequency (through drift or selection; Schaack *et al*. 2020) in order to meet the detection thresholds set by the high error rates (often above 10^-3^ errors per bp) of traditional DNA sequencing methods (Schirmer *et al*. 2016).

Two large-scale MA experiments with the nematode *Caenorhabditis elegans* have reached very different conclusions regarding the spectrum of mtDNA mutations (Denver *et al*. 2000; Konrad *et al*. 2017). The pioneering study of (Denver *et al*. 2000) was the first MA experiment to characterize mtDNA mutations in metazoans, using Sanger sequencing to detect a total of 26 mutations in 74 MA lines bottlenecked for an average of 214 generations. The 16 single nucleotide variants (SNVs) they identified indicated a bias towards mutations that increase GC content (10 variants increased GC content, four decreased GC content and two were GC neutral), a surprising finding given that the *C. elegans* mtDNA is 75.6% AT. However, a more recent *C. elegans* MA experiment utilized Illumina sequencing to detect a total of 24 mtDNA mutations (nine SNVs) in 20 MA lines each bottlenecked for an average of 363 generations and found a strong bias in the opposite direction; eight of nine SNVs increased AT content, and the other SNV was AT neutral (Konrad *et al*. 2017). Both of these studies were impressive, multi-year undertakings, and it is unclear whether the differences in the reported spectra are driven by biological differences in the *C. elegans* lines or rearing conditions, differences in sequencing techniques, or noise associated with small sample sizes. Additional MA experiments conducted with the nematodes *Caenorhabditis briggsae* (Howe *et al*. 2010; Wagner *et al*. 2020) and *Pristionchus pacifics* (Molnar *et al*. 2011) have also yielded very few mutations (seven to 19 SNVs), making it difficult to get precise estimates of mutation parameters.

An alternative to MA experiments has emerged in the form of high-fidelity sequencing techniques that can detect mtDNA variants segregating in tissues at extremely low frequencies (Salk *et al*. 2018; Sloan *et al*. 2018). One technique called Duplex Sequencing (Schmitt *et al*. 2012; Kennedy *et al*. 2014) is particularly useful for this application as it is highly accurate with error rates as low as ∼2×10^-8^ per bp (Wu *et al*. 2020), facilitating detection of *de novo* mtDNA mutations essentially as they occur. Duplex Sequencing works by tagging both ends of library molecules with random barcodes before they are amplified and sequenced. These barcodes are then used to create families of reads corresponding to each of the original strands from a parent DNA fragment. Consensus base calling eliminates variants present only in a minority of reads within a family or only in reads originating from one of the two strands, as such variants are common artifacts of single-stranded DNA damage, PCR misincorporations, and sequencing error. Here, we employed hybridization-based mtDNA enrichment coupled with Duplex Sequencing to greatly increase the detection of *de novo* mutations and better characterize the spectrum and distribution of mutations in the *C. elegans* mitochondrial genome.

## MATERIALS AND METHODS

### Nematode growth and DNA extraction

Replicate cultures of nematodes were derived from the Bristol N2 strain of *C. elegans* and grown on nematode growth media (NGM; He 2011) plates with *E. coli* strain OP50 at 20°C. Our nematode rearing protocol (outlined in Figure S1) was designed to 1) sample a diverse tissue pool comprised of many individuals (i.e. with a diversity of rare mtDNA mutations), 2) target lineages separated from each other for multiple generations to limit the contribution of shared, heteroplasmic mtDNA variants, and 3) limit the total number of generations (to three) in order to minimize potential biases of selection. Three sibling (F0) worms were randomly chosen from our lab stock of N2 nematodes to initiate experimental populations (referred to hereafter as populations 1, 2 and 3). Each population was maintained for 15 generations, with passages of ten adults at each generation, before a single adult F15 founder was randomly selected and transferred to a fresh NGM plate. Three F16 progeny were then randomly chosen from each population to initiate the nine replicates assayed in this study (referred to hereafter as replicates 1a, 1b, 1c, 2a, 2b, 2c, 3a, 3b, and 3c). Each replicate culture was allowed to proliferate for three generations (14 to 16 days) before all offspring in a replicate (mixed-stage individuals) were pooled for DNA extraction.

The three replicates from each population were grown in parallel, and the different populations were grown in three sequential batches. All culturing conditions and plate transfers followed the same outline (Figure S1). The replicates were checked daily to monitor growth and food supply. After each generation, worms were collected in M9 liquid buffer (He 2011), pelleted by centrifugation at 1,900 rcf for 30 sec, and redistributed onto fresh NGM plates. The first replating took place 7 to 9 days after the F16 replicate founder was plated. The F17 progeny were collected and redistributed onto 3 new plates. During this replating, one-third of the worms (by volume of the pellet) were discarded so as not to overwhelm the food supply on the new plates. 3 to 6 days later, the F18 worms were collected from the 3 plates (per each replicate) and pooled before half of the worms were discarded (again, so as not to overwhelm the food supply on the new plates), and the other half were redistributed onto 10 new NGM plates (per each replicate) for the final stage of growth. After 2 to 3 days, the F19 worms were collected from all 10 plates (per each replicate) and pooled into one 15 mL falcon tube (per each replicate). The worms were pelleted by centrifugation at 1,900 rcf for 30 sec, and the supernatant was discarded. Then, M9 buffer was used to resuspend the worms, and the wash process was repeated 2 more times to deplete contaminating *E. coli.* Total-cellular DNA was extracted from the worm pellet using the DNeasy Blood and Tissue Kit (Qiagen), following manufacturer’s instructions.

### Duplex library preparation and mtDNA enrichment

Total cellular DNA was stored at -20°C until all 9 replicates were processed. Then, Duplex Sequencing libraries were created for all 9 samples, following our previously described protocols (Wu *et al*. 2020) with some modifications. Briefly, DNA was fragmented with a Covaris M220 Focused-Ultrasonicator, end repaired (NEBNext End Repair Module), and A-tailed (Klenow Fragment Enzyme, 1mM dATP). The A-tailed DNA was then adaptor ligated with custom Duplex adaptors, which contain the 12-bp random barcodes necessary for double-stranded consensus building. The adaptor-ligated product was treated with a cocktail of three repair enzymes (Uracil-DNA Glycosylase, Fpg, and Endonuclease III in NEB CutSmart Buffer) to remove fragments with single-stranded damage and 16 ng of repaired DNA was used as input for the first round of PCR (13 cycles), in which Illumina adaptors with multiplexing indices were incorporated into the amplicons (NEBNext Ultra II Q5 Master Mix, custom IDT Ultramer primers).

Then, 336 ng of each amplified library was processed with the Arbor Biosciences myBaits Mito kit, following manufacturer’s instructions (manual version 4.01) and using their biotinylated bait panel specifically designed against the *C. elegans* mitochondrial genome. The enriched product was then PCR amplified for 11 additional cycles using universal p5 and p7 primers that anneal upstream of the multiplexing indices (NEBNext Ultra II Q5 Master Mix). Amplified libraries were separated and imaged on an Agilent TapeStation 2200 (High Sensitivity D1000 reagents) for quality control, and 80 nmol of each library was pooled for sequencing. The pooled library molecules had an average length of 369 bps.

### Total-cellular shotgun libraries to control for NUMT derived mapping artifacts

Rare variant detection in mitochondrial genomes is complicated by the fact that mitochondrial sequences can occasionally be transferred to the nuclear genome (Richly and Leister 2004). These mitochondrial derived nuclear genome sequences (referred to as NUMTs) accumulate point mutations as they are largely non-functional. NUMT-derived library fragments can map to the mitochondrial genome, and variants that have accumulated in NUMTs can be difficult to distinguish from true mitochondrial mutations (Hazkani-Covo *et al*. 2010). To overcome this challenge, we have previously implemented a *k*-mer count-based approach (Wu *et al*. 2020; Broz *et al*. 2021; Waneka *et al*. 2021), where counts of each putative mtDNA mutation are tabulated from a total-cellular shotgun DNA library. NUMT-derived variants are expected to have *k*-mer counts in the shotgun library that are similar to the counts for rest of the nuclear genome. In contrast, true mtDNA mutations are expected to have *k*-mer counts substantially lower (typically 0, barring sequencing errors and/or rare convergent mutations in the shotgun library) than the rest of the nuclear genome. We generated three replicate shotgun libraries for this purpose from total-cellular *C. elegans* DNA, following the same design used to generate the Duplex Sequencing replicates (Figure S1). Three sibling nematodes were used to initiate lineages that were propagated for three generations before nematodes from several plates (per replicate) were pooled for DNA extraction. To limit the potential contribution of shared, heteroplasmic variants in our nine replicates and the shotgun samples, the parent of the three replicates was selected from a lineage of N2 nematodes divergent from the nine replicates we assay for mtDNA mutations. DNA was extracted using the Qiagen DNA Blood and Tissue Kit following manufacturer’s instructions, and shotgun libraries were created using the NEBNext Ultra II FS DNA Library Prep Kit, with 50 ng of input DNA, a 15-minute fragmentation step, and 5 cycles of PCR amplification. Assessment of the shotgun libraries on the Agilent TapeStation 2200 (High Sensitivity D1000 reagents) revealed adaptor dimers, which were subsequently removed with size selection on a 2% BluePippin gel (Sage Science) using a target range of 300-700 bp. The resultant pooled sample had an average fragment length of 392 bp.

### Sequencing of shotgun and Duplex Sequencing libraries, variant detection and analysis

Total-cellular shotgun libraries were sequenced on an Illumina NovaSeq 6000 platform (2×150 bp reads) at the University of Colorado Cancer Center, resulting in 18.1 to 22.2 M read pairs per library, equating to roughly 60× coverage of the *C. elegans* nuclear genome. The Duplex Sequencing libraries were sequenced on an Illumina HiSeq4000 platform (2×150 bp reads) by Novogene in two runs, resulting in a total of 66.8 to 76.9 M read pairs per library.

Duplex Sequencing reads were processed with our previously described pipeline (https://github.com/dbsloan/duplexseq; Wu *et al*. 2020), which 1) trims adaptor sequences (cutadapt v1.16; Martin 2011), 2) calls duplex consensus sequences (DCSs) based on shared random barcodes (each DCS required a minimum of six raw Illumina reads – at least three from each strand), 3) filters out discordant DCSs with ambiguous bases resulting from disagreement between strands, 4) aligns concordant DCSs to the reference genome (NCBI reference sequence NC_001328.1; bowtie2 v2.3.5; Langmead and Salzberg 2012), 5) calls variants and DCS coverage by parsing the DCS alignment and 6) filters NUMTS through a comparison to a *k*-mer database generated from the shotgun libraries (KMC v 3.0.0; https://github.com/refresh-bio/KMC). An expected AT equilibrium was calculated as:

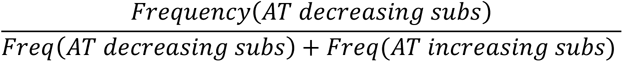

To account for the nested structure of our data (i.e. replicates a, b, and c are nested within each population 1, 2, and 3), we implemented mixed linear models in R (ver 1.3.959), using the lme4 package. In such analyses, the genomic comparison of interest (for example substitution class or genome region) was set as a fixed effect, and replicate was set as a random effect nested within population, which was also set as a random effect.

## RESULTS AND DISCUSSION

### Targeted Duplex Sequencing provides a high-resolution view of mutation in the *C. elegans* mitochondrial genome

We enriched for mtDNA derived sequences through hybrid capture, which was highly effective, as 99.91% of all DCSs mapped to the *C. elegans* mitochondrial genome (each DCS is a consensus sequence produced from at least 6 concordant Illumina reads). In contrast, in a preliminary trial with Duplex Sequencing libraries made from total cellular *C. elegans* DNA (SRR14352239), less than 0.5% of DCSs mapped to the mitochondrial genome. For each of the nine replicates assayed, the average DCS coverage of the mitochondrial genome was 13,257×, with a range from 9,099× in replicate 1b to 15,451× in replicate 1a (Table S1).

We identified a total of 456 DCSs with single nucleotide variants (SNVs) and 17,582 DCSs with indels (SNVs: File S1, indels: File S2). These counts do not include one variant identified as a NUMT artifact based on *k*-mer counts in total-cellular shotgun libraries. The putative NUMT was present at a low frequency in every replicate and mapped to the only NUMT to have been previously identified in the *C. elegans* nuclear genome (Frith 2011). The above counts were tabulated after correcting three positions with fixed or nearly fixed differences in the nine replicates (Table S2), which presumably represent existing differences in our lab N2 line compared to the published N2 mitochondrial genome (NC_001328.1). Raw DCS counts are inflated however by several variants present at high enough frequency to be detected in numerous DCSs. For example, a CG→TA mutation at position 5079 was present at a frequency of 0.004 in replicate 1c (158 DCSs). Similarly, a 1-bp A insertion at position 3235 was present at a frequency of 0.52 to 0.65 in the population 2 replicates (>15,000 DCSs). Across all replicates, there were 234 unique sites with an SNV, 36 unique sites with an insertion, and 65 unique sites with a deletion. Although most SNVs were “singletons” detected in only one DCS, we did identify 23 SNVs that were represented by multiple DCSs from the same replicate.

Interestingly, there were also 15 sites with SNVs shared between replicates from different populations. Mutations shared between replicates could have either arisen through two independent, convergent mutations, or the mutation could have occurred just once in the common ancestor of both replicates and remained heteroplasmic in both replicates. Given that the majority of these SNVs are shared across populations separated for 15 generations before replicate subdivision (File S1; Figure S1), we posit that most shared SNVs arose through independent mutations. This conclusion is further supported by the observation that no SNVs were exclusively shared among replicates from the same population (File S1). Therefore, for all downstream analyses we interpreted shared variants as independent counts, resulting in 253 SNVs, 108 insertions, and 84 deletions (SNVs: File S1, indels: File S2). The 253 SNVs include six sites which were tri-allelic. In all six cases, a CG reference base pair experienced both a CG→TA transition and a CG→AT transversion. Three of these six sites were tri-allelic within a single replicate. We observed only a single dinucleotide substitution: an AA→GG change at position 5010 in replicate 1c. The count of 445 total mutations (253 SNVs plus 192 indels) observed here is 9-fold higher than the highest count (51 mutations; Howe *et al*. 2010) detected in any nematode MA experiment to date (Denver *et al*. 2000; Molnar *et al*. 2011; Konrad *et al*. 2017; Wagner *et al*. 2020). Although this bulked sequencing approach of whole nematode populations is not conducive to estimating an absolute mutation rate per generation, the large number of detected variants makes it a powerful one for characterizing the spectrum and distribution of mutations.

### Low-frequency variants detected in *C. elegans* mtDNA indicate a strong bias towards mutations that increase AT content

We found significant variation in the (log transformed) SNV frequency between the six substitution classes (one-way ANOVA, *p* << 0.0001; Figure 1a). CG→TA transition and CG→AT transversion frequencies were similar to one another (2.09×10^-7^ and 1.81×10^-7^, respectively), and 3- to 16-fold higher than the frequencies in the other substitution classes. Previous MA experiments to analyze nematode mtDNA mutation spectra have also reported a dominance of CG→TA transitions (Howe *et al*. 2010; Konrad *et al*. 2017), but only one *C. briggsae* study (with a count of 10 total SNVs), reported CG→AT transversions at a similar relative abundance to CG→TA transitions (Wagner *et al*. 2020).

**Figure 1.**
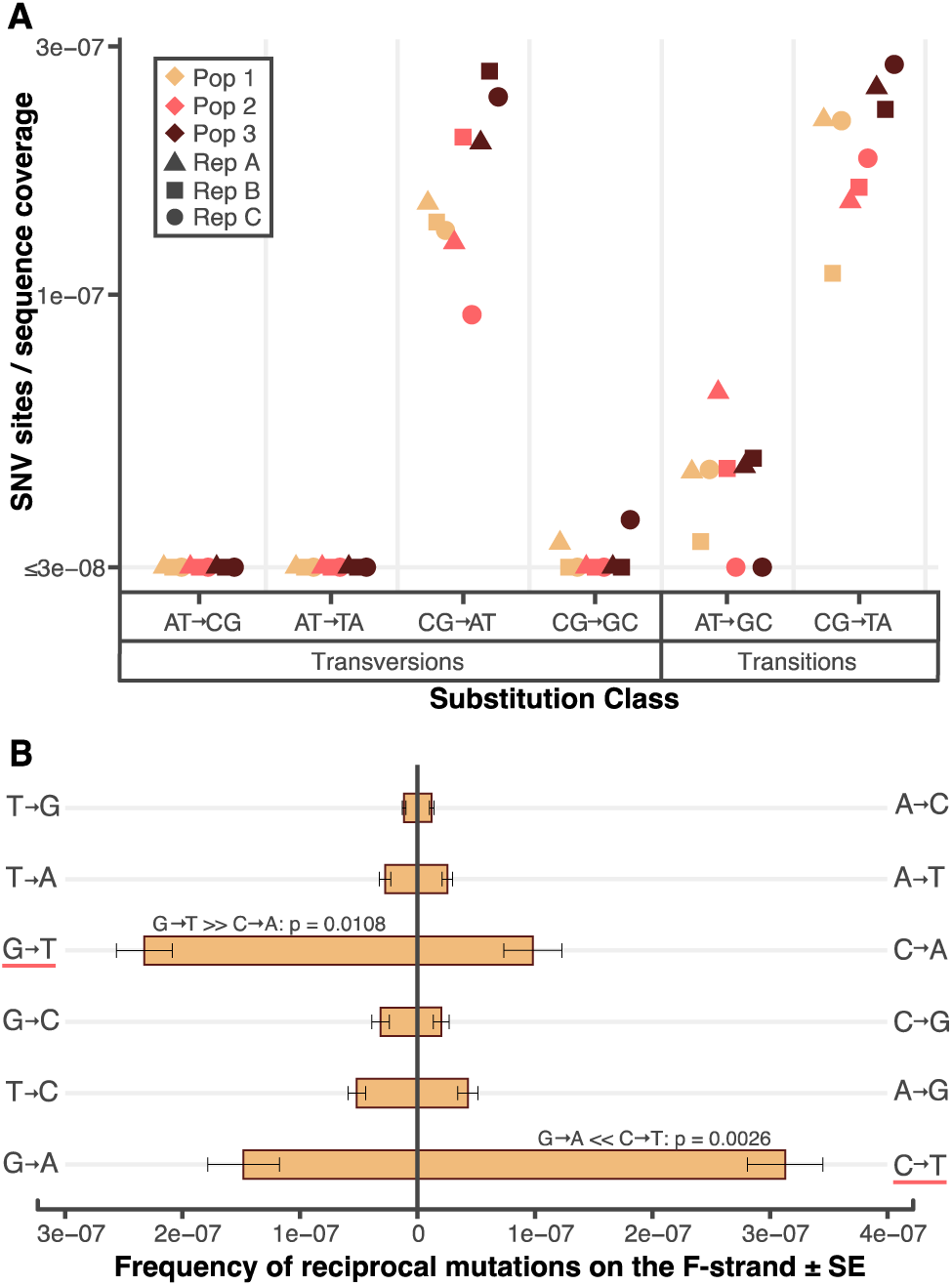
The *C. elegans* mtDNA mutation spectrum is dominated by mutations that increase AT content and exhibits strand asymmetry. **(A)** Variation in the frequency of mutations across six substitution classes. CG→AT transversions and CG→TA transitions are the most abundant substitution types. Each point on the plot represents one of the nine replicates assayed. SNV frequencies were calculated as the number of sites in a replicate with a mutation normalized by the coverage of the corresponding base pair type. For example, the CG→AT mutation frequency shows the CG→AT event count divided by GC coverage for each replicate. A floor was applied to frequencies below 3e-08, which approaches the error threshold of duplex sequencing (Wu *et al*. 2020). **(B)** Strand asymmetry of mutations in *C. elegans* mtDNA. Both CG→AT and CG→TA substitutions show significant strand asymmetry (one-way ANOVA, *p*-values noted on figure), with the G→T and C→T changes (underlined in red) occurring predominately on the forward (F) strand, which in *C. elegans* mtDNA is synonymous with the heavy-strand and for genic regions also the coding-strand. Mutation frequencies were calculated as the average of the nine replicates and were normalized by the sequencing coverage of each base type on the F- strand. For example, the G→T mutation frequency shows all G→T events on the F-strand divided by G coverage on the F-strand, and the C mutation frequency shows all C events on the F-strand divided by C coverage on the F-strand.

We asked if the *C. elegans* mtDNA is at AT equilibrium, in which case the number of AT-decreasing mutations would equal the number of AT-increasing mutations. Given the dominance of CG→TA transitions and CG→AT transversions in the spectrum (Figure 1a), it was not surprising that the AT-increasing count of 173 was significantly greater than the AT- decreasing count of 52 (binomial test, *p* << 0.0001). This difference is not driven by differences in AT vs. GC sequencing coverage, as the coverage-adjusted AT-decreasing count (normalized by the ratio of AT coverage per base pair over GC coverage per base pair, as in Waneka *et al*. 2021) of 61 was still significantly less than the adjusted AT-increasing count of 164 (binomial test, *p* << 0.0001). We used the coverage-adjusted counts to determine an expected AT equilibrium of 89.4% (formula in Methods). This expected value is substantially larger than the actual mtDNA AT content (75.6%), slightly greater than the AT content at 4-fold degenerate sites (86.4%) and slightly less than AT content of the two intergenic regions of the *C. elegans* mtDNA (90.8% AT). These results suggest that AT mutation bias would push the *C. elegans* mtDNA to even more extreme AT contents if not for the stabilizing effects of natural selection, thus supporting the findings of Konrad *et al*. (2017) which ran contrary to the earlier report of a mutational bias towards GC in *C. elegans* mtDNA (Denver *et al*. 2000).

### Asymmetries between forward and reverse mtDNA strands suggest differences in single-stranded damage

Several biological processes, including replication (Kennedy *et al*. 2013) and transcription (Liu and Zhang 2020), can lead to systematic differences in the amount of DNA damage experienced by the two DNA strands. By definition, mutations affect both strands of DNA, but single-stranded asymmetries can be studied by comparing the frequency of reciprocal single-stranded changes for each substitution class (for example C→T vs. G→A changes) on a given DNA strand. The coding sequence for all of *C. elegans* mtDNA genes (12 protein coding genes, the two rRNA genes, and the 22 tRNA genes) are oriented in the same direction on one DNA strand (hereafter F-strand for forward strand) (Okimoto *et al*. 1992). To search for signatures of DNA damage in our Duplex Sequencing data, we performed a strand asymmetry test for each of the six substitution classes. This analysis revealed significant asymmetries in both CG→TA transitions and CG→AT transversions (one-way ANOVAs, *p* = 0.0026 and 0.0108, respectively), with disproportionate amounts of C→T and G→T changes occurring on the F-strand (Figure 1b).

### The role of oxidative damage in the *C. elegans* mtDNA mutation spectrum and strand asymmetries

CG→AT transversions are indicative of oxidative damage because oxidized guanines (e.g. 8- oxo-G) are often mis-paired with adenine, causing G→T changes (Kennedy *et al*. 2013; Kino *et al*. 2017). The relative abundance of CG→AT transversions in the *C. elegans* mtDNA Duplex Sequencing spectrum (Figure 1a) compared to mtDNA mutation spectra produced through high-fidelity sequencing in other metazoans (Kennedy *et al*. 2013; Itsara *et al*. 2014; Ni *et al*. 2015; Samstag *et al*. 2018; Arbeithuber *et al*. 2020) suggests that oxidative damage may be particularly important for driving mtDNA mutation in nematodes. The 2.3-fold enrichment of G→T changes on the F-strand (Figure 1b) provides evidence that the abundance of CG→AT transversions observed here is not artifactual and suggests the F-strand suffers increased loads of oxidative damage *in vivo*. The other common substitution class in our data, CG→TA transitions, are likely also driven by strand-specific damage (cytosine deamination), given the 2.1-fold enrichment of C→T changes on the F-strand. Cytosine deamination can be related to oxidative damage, but it can also occur spontaneously via hydrolysis in the absence of oxidative damage (Kreutzer and Essigmann 1998; Nabel *et al*. 2012).

Previous high-fidelity sequencing studies observing C→T vs. G→A strand asymmetries in mtDNAs of fruit flies (Itsara *et al*. 2014), mice (Arbeithuber *et al*. 2020) and humans (Kennedy *et al*. 2013) have hypothesized that increased spontaneous deamination of cytosine on the F-strand (referred to as the H-strand or minor-strand in those systems) occurs due to increased single-stranded exposure during mtDNA replication (Falkenberg 2018). *C. elegans* mtDNA replication is distinct from the theta-type mtDNA replication that has been heavily characterized in vertebrates (Yasukawa *et al*. 2006; Cluett *et al*. 2018; Falkenberg 2018) and has also been documented in fruit flies (Jõers and Jacobs 2013). In *C. elegans*, mtDNAs are replicated through a rolling circle mechanism which produces double-stranded concatemers up to 48.2 kb in length (3.5× the length of a single mtDNA; Lewis *et al*. 2015). In characterizing the rolling circle mechanism of mtDNA replication utilized by *C. elegans,* Lewis *et al*. (2015) found some evidence of single-stranded DNA through transmission electron microscopy but noted that “replication intermediates lack the extensive single-stranded DNA character expected” from theta-type mtDNA replication. Interestingly, the magnitude of asymmetries observed here (2.3- fold for G→T changes and 2.1-fold for C→T changes on the F-strand) are substantially lower than the C→T enrichment on equivalent strands in aged mice (7.9- to 11.9-fold depending on the tissue-type; Arbeithuber *et al*. 2020), humans (Kennedy *et al*. 2013), and fruit flies (Itsara *et al*. 2014). In the latter two studies, the magnitudes of strand asymmetries were not reported explicitly but were >10-fold based on study data. We posit that the reduced magnitude of strand asymmetries in *C. elegans* may be associated with the lack of single-stranded intermediates reported by Lewis et. al (2015). Transcription may also drive mutational asymmetries observed here, as the coding (or sense) strand may be exposed while RNA polymerases bind and read off of the template strand (Liu and Zhang 2020). Because the template sequences for all *C. elegans* mtDNA genes are located on the same strand, it is unclear if the F-strand suffers increased single-stranded exposure due to replication, transcription or both.

Interestingly, rolling circle replication can apparently be induced in mtDNAs of human cells through treatment with H_2_O_2_ (a ROS; Ling *et al*. 2016; Ling and Yoshida 2020), indirectly supporting the link between rolling circle replication and oxidative stress in *C. elegans* mtDNAs. Assays with human Pol γ *in vitro* reveal the polymerase is particularly prone to misincorporations leading to AT→GC and CG→TA transitions (Longley *et al*. 2001; Zheng *et al*. 2006). Misincorporations by the nematode Pol γ have not been characterized, so it is possible that the relative abundance of CG→AT transversions in the Duplex Sequencing spectrum could reflect distinct replication errors of the *C. elegans* Pol γ. However, such polymerase error would be unable to explain the G→T vs. C→A strand asymmetries that we observed (Figure 1b).

In metazoan mtDNAs, BER of oxidized guanines is hypothesized to be mediated by mitochondrially targeted glycosylases OGG1 and/or MUTYH. However, *ogg1* mutant flies (Itsara *et al*. 2014) and *ogg1*/*mutyh* double mutant mice (Kauppila *et al*. 2018) show no increase in mtDNA mutations compared to wild-type individuals, even when mitochondria ROS levels are elevated through knockout of the mitochondrially targeted superoxide dismutase (*Sod2)* in these lines. Interestingly, *C. elegans* lines lacking mitochondrially targeted SODs experience significantly elevated mtDNA damage compared to wild-type lines, as measured with short- and long-amplicon quantitative real-time PCR (Ng *et al*. 2019). mtDNA mutations have not yet been assessed in *C. elegans sod* mutants or in lines with deficiencies in BER. While the relative abundance of CG→AT transversions in our Duplex Sequencing data supports a role of oxidative damage in driving *C. elegans* mtDNA mutations, we do not consider this evidence in support of the mitochondrial free radical theory of aging (mFRTA). The mFRTA posits that oxidative stress is causal to aging (Harman 1972). While we find evidence that oxidative stress may be causal to mtDNA mutations in *C. elegans*, previous studies which have more explicitly tested the mFRTA in *C. elegans* have not found a consistent link between oxidative stress and nematode lifespan (Gruber *et al*. 2011; Ng *et al*. 2019).

### Comparisons to other mtDNA mutation studies

It is unlikely that the relative abundance of CG→AT transversions reported here is an artifact of Duplex Sequencing because we have recovered a diversity of unique mutation spectra with this same technique in various other biological systems (Wu *et al*. 2020; Broz *et al*. 2021; Waneka *et al*. 2021), some of which have shown very low relative frequencies of CG→AT transversions. Further, Duplex Sequencing, when used to measure mutations in human and mice mtDNAs, yielded spectra with very few CG→AT transversions (Kennedy *et al*. 2013 and Arbeithuber *et al*. 2020, respectively). Different high fidelity techniques have also revealed spectra relatively deplete of CG→AT transversions in the mtDNAs of wild-type fruit flies (Itsara *et al*. 2014) and mice (Ni *et al*. 2015).

Given that we detected mutations in pooled somatic and germline tissues, we considered the possibility that the relative abundance of CG→AT transversions in the Duplex Sequencing spectrum (Figure 1a) compared to what has been reported in nematode MA studies (Howe *et al*. 2010; Molnar *et al*. 2011; Konrad *et al*. 2017) could reflect distinct mutational spectra in somatic vs. germline mtDNAs. Such a distinction would imply that oxidative damage and associated CG→AT transversions are more prevalent in somatic mtDNAs than in the mtDNAs maintained in the nematode germline. The mixed-stage populations from which we extracted DNA likely included some older individuals (no older than 14-16 days old, the total time of replicate growth; Figure S1), potentially increasing the contribution of somatic mtDNA mutations (Kennedy *et al*. 2013; Arbeithuber *et al*. 2020). However, studies of mtDNA replication across *C. elegans* development in mutant lines with deficiencies in Pol γ have established that mtDNA replication occurs primarily in the nematode gonad, such that essentially all somatic mtDNAs originate at embryogenesis (Bratic *et al*. 2009). The embryonic origin of somatic mtDNAs therefore blurs the distinction between somatic and germline mtDNAs in nematodes. If the abundance of CG→AT transversions is caused by oxidative damage to mtDNA in somatic tissues, it would indicate that there is some degree of active mtDNA replication or erroneous mtDNA repair converting single-stranded DNA damage into double-stranded mutations detectable with Duplex Sequencing.

Given that our mitochondrial DCS coverage (13,257× on average) is well below the estimated number of nematodes in each tissue pool (∼50,000 assuming each of the 10 plates pooled for each replicate contained ∼5,000 nematodes), we have likely sampled less than one mtDNA per nematode. Therefore, the 23 mutations that were detected in > 1 DCS in our dataset are more likely to be inherited, germline mtDNA mutations, present in multiple individuals within a replicate. Importantly, these sites yielded a spectrum with a similar frequency of CG→AT transversions (35% of all substitutions; Table S3) as in the full dataset (32% of all substitutions; Table S3; Figure 1a). Still, it is possible that the spectrum reported here is influenced in part by distinct mutational patterns in somatic mtDNAs.

Another potential explanation for the differences in the Duplex Sequencing results compared to those from MA studies is that our spectrum consists mostly of extremely rare variants captured by only a single DCS (230 of 253 observed SNVs), whereas nematode MA studies have applied detection cutoffs requiring variants to be present in at least 3 or 4 unique reads (for Illumina based studies with coverages of approximately 388 or 282; Konrad *et al*. 2017 and Wagner *et al*. 2020, respectively) or simply as ‘detectable on a chromatogram’ for Sanger based studies (Denver *et al*. 2000; Howe *et al*. 2010; Molnar *et al*. 2011). Given that the mutations detected in MA studies were initially generated as only a single copy within an individual and had to rise in frequency to meet detection thresholds, it seems likely that the MA mutation spectra could be biased by selection. Even small selective biases may have dramatic effects on observed spectra given that *C. elegans* MA studies passaged lineages for 214 or 363 generations (roughly two or three years of propagation; Denver *et al*. 2000 and Konrad *et al*. 2017, respectively), whereas our culturing design allowed just three generations for mutations to occur (Figure S1). By minimizing the number of generations, we may have reduced the opportunity for within-individual selection to shape the mutation spectrum, although this trades off with the fact that the absence of the bottlenecking used in MA lines would allow for selection to act at an organismal level in these three generations.

If CG→AT transversions experience stronger negative selection than CG→TA transitions, which is plausible since transversions are more likely to result in amino acid changes (Okimoto *et al*. 1992), they could be underrepresented in MA studies due to within-individual selection pressures. Indeed, in our Duplex Sequencing Data 93.4% (57/61) of CG→AT transversions but only 75.6% (61/82) of CG→TA transitions result in amino acid changes (File S1). This same logic has been used to explain an underrepresentation of nuclear CG→AT transversions in natural nematode populations compared to the relative abundance of nuclear CG→AT transversions in multiple nematode MA studies (Denver *et al*. 2012; Weller *et al*. 2014) because in that comparison, the natural population spectrum is expected to be more strongly affected by selection (Rajaei *et al*. 2021). Konrad et al (2017) compared *C. elegans* mtDNAs from 38 natural isolates (sequenced in Thompson *et al*. 2013) and found that transitions (including both CG→TA and TA→CG changes, which cannot be reliably polarized in the population dataset) account for 83% of the 408 observed substitutions, yielding a transition/transversion ratio (hereafter Ti/Tv ratio) of 4.75 (Table S4). The dominance of transitions at the population level appears to be driven in part by stronger selection against transversions because constraining the population data set to the 162 substitutions at four-fold degenerate sites yields a reduced Ti/Tv ratio of 3.26 (Table S4). The four-fold degenerate Ti/Tv ratio is still substantially higher that the Ti/Tv ratio we observe with Duplex Sequencing (1.32; Table S4). Considering that four-fold degenerate sites are expected to experience minimal selection, the elevated Ti/Tv ratio at four-fold sites compared to the one from our Duplex Sequencing data suggests the latter may not be fully representative of inherited mtDNA mutations in natural *C. elegans* populations.

### Distribution of mtDNA SNVs across the mitochondrial genome

The large number of SNVs we detected with Duplex Sequencing allowed us to study how these events are distributed across the genome. As shown in the middle track (yellow histogram) of Figure 2a, the depth of DCS coverage varied substantially across the genome. Much of this variation can likely be attributed to differences in local GC content, as the AT DCS coverage (summed across replicates) accounted for only 72.1% of all DCS coverage, despite the fact that the mitochondrial genome is 75.6% AT. The AT vs. GC coverage disparity is exaggerated in regions with long stretches of sequence that are AT rich, as the 10% of 50 bp windows with the lowest GC content have 8.9-fold lower DCS coverage than windows of median GC content, and 17.5-fold lower DCS coverage than the 10% windows with the highest GC content (Figure S2). Still, 95.6% of 50-bp windows had DCS coverage (summed across nine replicates) above 1000×. Bias against AT-rich sequences during library amplification and decreased binding affinities for AT rich baits during mtDNA enrichment both likely contribute to decreased AT coverage.

**Figure 2.**
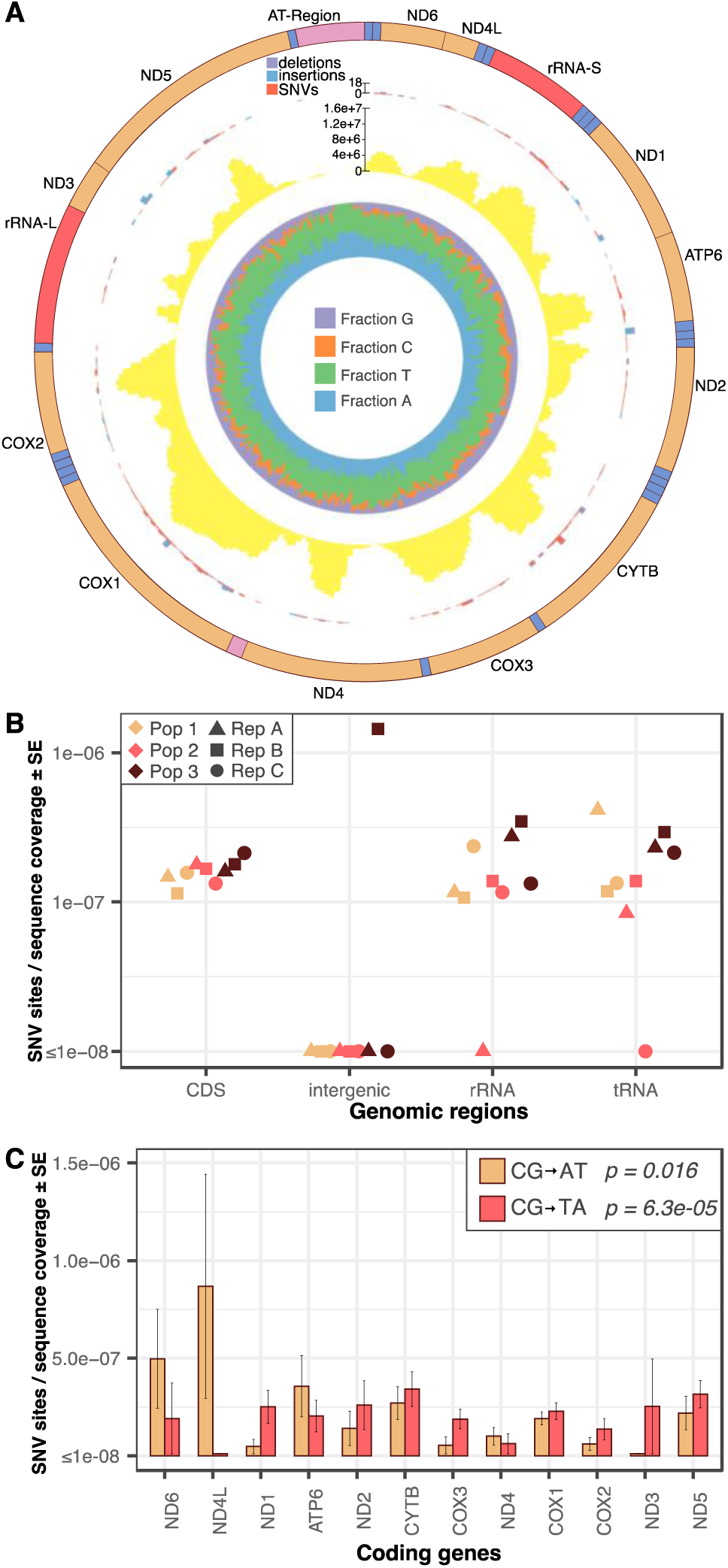
The distribution of mutations across the mitochondrial genome. (A) Map of the *C. elegans* mtDNA and summary of Duplex Sequencing data. The outermost track depicts the gene order and type, with protein-coding (CDS) genes shown in tan, rRNA genes shown in red, tRNA genes shown in blue, and intergenic regions shown in pink. The next track in from the outside depicts the total mutation counts in 50-bp windows, with deletions, insertions and SNVs colored differently according to the key at the top of the track. The next track in from the outside (yellow histogram) depicts the cumulative (sum of nine replicates) DCS coverage in 50-bp windows, with a scale bar included at the top of the track. The innermost track shows the relative fraction of each base type in 50-bp windows, with colors specified by the key in the figure center. The figure was generated with Circos v0.69-8 (Krzywinski *et al*. 2009). **(B)** Variation in SNV frequencies by genomic region. Note that low intergenic coverage resulted in the detection of only a single intergenic substitution (in replicate 3b). The other replicates had intergenic substitution counts of zero, but also had extremely low intergenic coverage, making comparisons that include intergenic SNV frequency low powered (see main text). No significant variation is observed among CDS, tRNA, or rRNA genes when the intergenic region is excluded. **(C)** Significant variation in SNV frequencies across the 12 protein coding genes was observed for only the two most common substitution classes (CG→AT transversions and CG→TA transitions). See Figure S3 for the gene specific mutation frequencies of all substitution classes.

After correcting for differences in coverage, we found no variation in mutation frequency between intergenic regions, protein coding genes, rRNA genes or tRNA genes (one-way ANOVA, *p =* 0.99; Figure 2b). However, comparisons against intergenic sequences are low powered given the relative lack of intergenic coverage (the largest of the two non-coding regions is extremely AT rich: 465 bp, 93.3% AT). A previous Duplex Sequencing study also found no variation across intergenic regions, protein coding genes, rRNA genes or tRNA genes in both wildtype fruit flies and in lines with a proofreading-deficient Pol γ (Samstag *et al*. 2018). We then tested for differences in mutation rates among protein coding genes, which comprise 74.5% of the *C. elegans* mtDNA. We found significant variation (one-way ANOVA, *p* = 0.0072; Figure S3) driven by between-gene differences in CG→TA transition and CG→AT transversion frequencies (one-way ANOVAs, *p* = 0.016 and 6.3×10^-5^ respectively; Figure 2c).

The cause of differences in SNV frequencies between genes remains unclear. Given the aforementioned mutational bias away from GC base pairs, we considered if differences in GC content among genes could be driving differences in SNV frequencies. There is a weak, positive correlation between gene-wide GC content and gene specific CG→TA transition frequencies (Pearson correlation: r=0.32, *p*=0.304; Figure S4) and a negative correlation between GC content and gene specific CG→TA transversion frequencies (Pearson correlation: r=-0.51, *p*=0.086; Figure S4), but neither was significant. Therefore, GC content variation does not explain differences in gene-specific mutation frequencies.

CG→TA transitions and CG→AT transversions both also show significant strand asymmetries (Figure 1b), so it seems possible that distance from an origin of replication (Kono *et al*. 2018) or differential transcription (Gaillard and Aguilera 2016; Wang *et al*. 2016) could play a role in driving mutation rate differences among genes (Figure 2c). The intergenic region upstream of *ND6* (labelled ‘AT region’ and drawn in pink in Figure 2a) contains short repetitive elements which led several to propose this region may be analogous to the D-loop in mammalian mtDNAs, acting as the F-strand replication origin (Lemire 2005; Bratic *et al*. 2010). However, this region lacks a GC skew inflection point typical of replication origins (Kono *et al*. 2018) and fails to form bubble arcs indicative of replication origins in two-dimensional gel electrophoreses (Lewis *et al*. 2015). Given the lack of evidence surrounding the location of a replication origin in the *C. elegans* mtDNA, it is difficult to assess how distance from replication origins may be impacting among-gene mutation rate differences. It is unlikely that differences in expression drive among gene mutation rate differences, as the *C. elegans* mtDNA is likely transcribed as a polycistronic RNA (Blumberg *et al*. 2017), with differences in relative abundances of specific mRNA transcripts presumed to arise from differences in mRNA stability and decay (D’Souza and Minczuk 2018). Reverse-transcriptase droplet digital PCR (ddPCR) estimates reveal relatively small differences (∼9-fold) in mRNAs levels among protein coding genes in the *C. elegans* mitochondria, while rRNAs are 50 to 200-fold more abundant than mRNAs (Held and Patel 2020). If transcription does drive mutation *C. elegans* mtDNAs, our finding of no difference in mutation rates between rRNA coding genes and protein coding genes (Figure 2b) supports the hypothesis that different levels of rRNA vs. mRNA arise through increased rRNA stability or mRNA decay (Held and Patel 2020). Yet another possibility, discussed below, is that differences in mutation frequencies among genes could be driven by local sequence features that are correlated with mutation and vary among genes.

### Distribution of mtDNA SNVs based on local sequence variation

Previous analyses of mtDNA mutations in flies (Samstag *et al*. 2018) and mice (Ni *et al*. 2015) have shown that the identities of neighboring nucleotides can have large impacts on mtDNA mutation frequencies. To understand if local sequence contexts influence SNV frequencies in the *C. elegans* mtDNA, we compared the variant frequencies at the 16 trinucleotide contexts (i.e., the mutated site and the flanking 5′ and 3′ nucleotides) for each substitution class and found significant effects for both CG→TA transitions and CG→AT transversions (one-way ANOVAs, *p =* 0.040 and 3.6×10^-5^, respectively), but not in any of the other substitution classes (Figure 3). It is likely that CG→TA transitions and CG→AT transversions are the only substitution types to show significant trinucleotide variation because they make up the majority of detected SNVs, while tests for variation in the other substitution classes are comparatively low powered. Different trinucleotides are apparently important in the two significant substitution classes. CG→TA transitions are particularly common at GCC/GGC, GCG/CGC and CCC/GGG trinucleotides (written 5′ to 3′). In contrast, CG→AT transversions are particularly common at ACT/AGT and ACG/CGT trinucleotides. It is possible that the above reported genic mutation rate variation among protein coding genes (Figure 2c) may be driven by a nonrandom distribution of ‘mutagenic’ trinucleotides.

**Figure 3.**
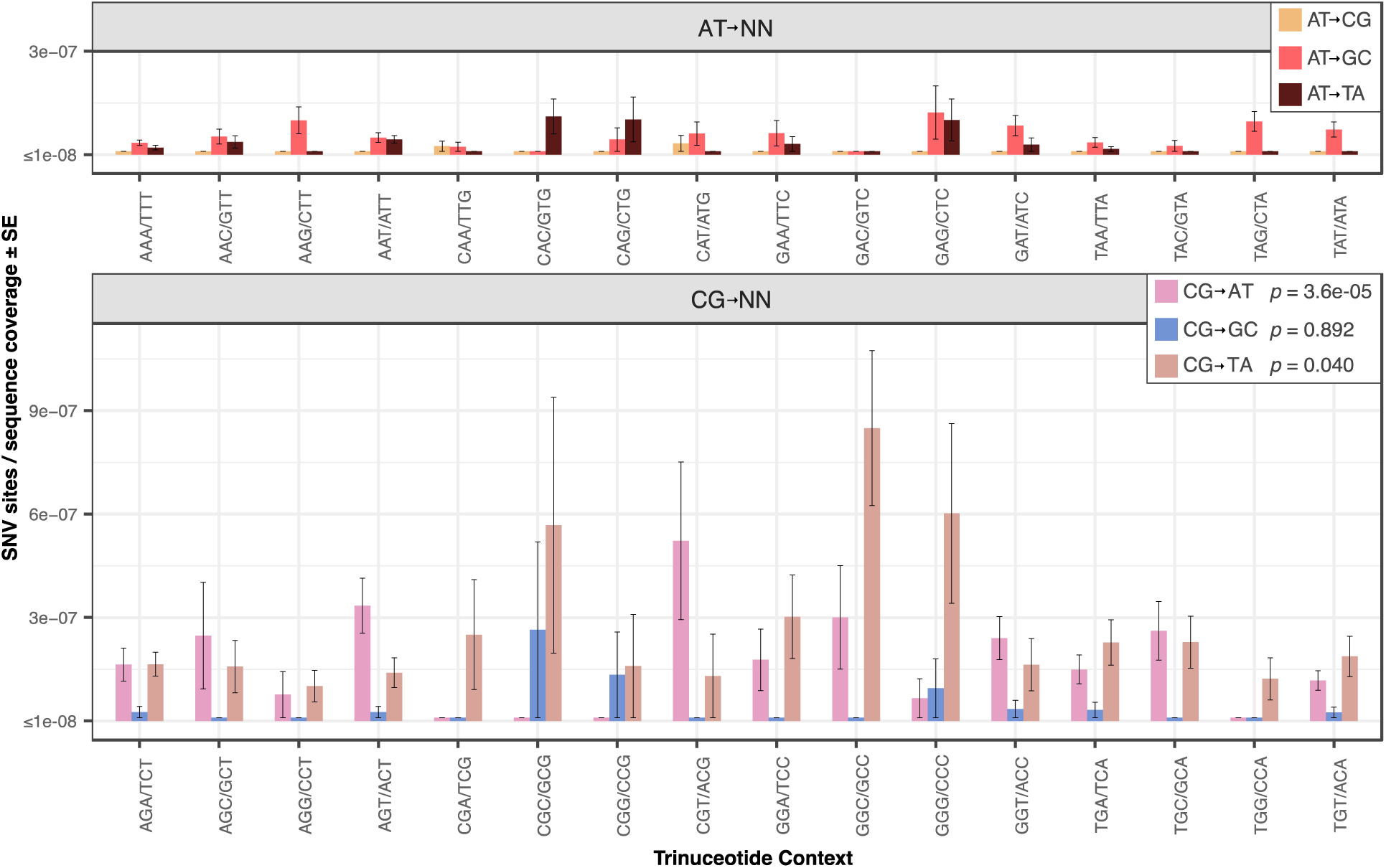
Variation in AT→NN (top panel) and CG→NN (bottom panel) mutation frequency across different trinucleotide contexts, where NN refers to any other base-pair. Significant variation was seen in the trinucleotide contexts for CG→AT transversions and CG→TA transitions (one-way ANOVA, *p*-values in figure legend).

### Nonsynonymous mutations are slightly more abundant than predicted by neutral simulations

To assess whether the identified sample of mutations was biased by selection, we used a simulation-based approach to obtain a neutral expectation for the ratio of nonsynonymous to synonymous (NS:S) mutations. There were 205 observed SNVs in protein-coding sequences (File S1), which we simulated onto a concatenation of the protein coding sequence. We attempted to control for the *C. elegans* mutation spectra and probability of detection by simulating the same number and type of each substitution from our observed data. This simulation was repeated 10,000 times to obtain a distribution of NS:S ratios. Our observed NS:S ratio of 3.01 was 1.3-fold higher than the median simulated value of 2.23 (*p =* 0.075). This result suggests that neither synonymous nor nonsynonymous substitutions are significantly overrepresented in the observed Duplex Sequencing dataset (Figure 4). As such, there is no evidence that purifying selection has played a large role in filtering this pool of low-frequency variants. Why the observed ratio contained a (marginally) higher proportion of nonsynonymous substitutions than the simulated ratio is not clear, though a previous study of mtDNA mutations in *Drosophila melanogaster* reported a significant overabundance of nonsynonymous substitutions compared to neutral expectations in a similar simulation based test (Samstag *et al*. 2018). Those authors proposed that deleterious mutations may reduce mitochondrial function, thus reducing the potential for oxidative damage and by extension make mutant bearing mitochondria “less prone to targeted degradation by quality control surveillance” (Samstag *et al*. 2018).

**Figure 4.**
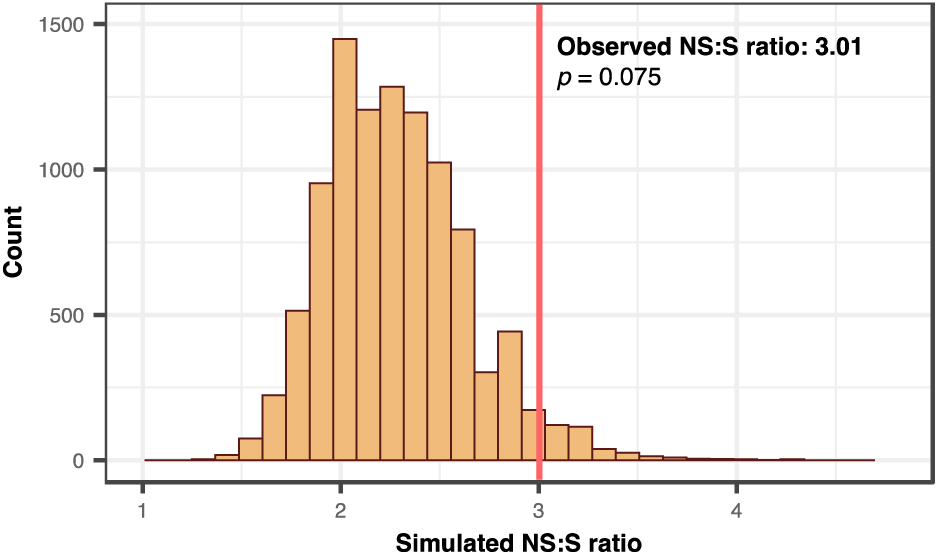
Simulation of mutations to derive null expectation of the ratio of nonsynonymous: synonymous substitutions (NS:S). The observed ratio of NS:S (marked with the vertical red line) is 1.3-fold higher than the null expectation generated from 10,000 simulations where the observed number and type of coding sequence mutations were randomly mapped onto the coding sequence, though this difference is not significant. The two-tailed *p*-value was derived by dividing twice the number of simulated NS:S ratios with values greater than the observed NS:S ratio by the total number of simulations (378*2/10000).

### Indels are less common than SNVs and are predominantly expansions or contractions of existing homopolymers

The aforementioned *C. elegans* MA experiments reached different conclusions regarding the relative abundance of indels vs. SNVs in mtDNAs, with the original study reporting a indel:SNV ratio of 0.65 (Denver *et al*. 2000) and the later study reporting a ratio of 1.42 (Konrad *et al*. 2017). As noted above, we found that many identical indels were shared across replicates and inferred that these shared variants arose from independent events, though we cannot rule out the possibility that they are ancestral, heteroplasmic variants. Assuming shared variants are independent events, we calculate an indel:SNV ratio of 0.75, supporting the finding that SNVs are more common than indels in the *C. elegans* mtDNA (Denver *et al*. 2000). This conclusion holds if we treat shared mutations as the products of a single event (shared ancestry), giving an indel:SNV ratio of 0.43. Duplex Sequencing coupled with hybridization-based enrichment is not designed to detect large deletions, which have been shown to accumulate at a high rate in mtDNAs of *C. briggsae* (Howe *et al*. 2010; Wagner *et al*. 2020), so we may be underestimating the total number of mtDNA indels. If indels are more prone to homoplasy *within* replicates (than SNVs), which may well be the case at long homopolymers (single-nucleotide repeats), this would lead us to further underestimate the indel:SNV ratio.

We find different results for the relative abundance of insertions vs. deletions depending on how we categorize shared mutations. Deletions are about ∼2-fold more abundant than insertions (65 vs. 36, respectively) when we assume that shared mutations arose through common ancestry. However, when assuming shared indels are independent events that arise through homoplasy, we see a difference in the other direction (84 deletions vs. 108 insertions). This discrepancy reflects the fact that identical insertions were more commonly shared between replicates than were identical deletions. We surveyed the sequence surrounding each of the observed indels and found that the majority of both insertions and deletions are 1-bp expansions or contractions of existing homopolymers, which is consistent with previous *C. elegans* MA studies (Denver *et al*. 2000; Konrad *et al*. 2017). This pattern holds regardless of whether we assume shared indels arose through homoplasy (Figure 5) or common ancestry (Figure S5). Accordingly, the length of both insertions and deletions skews heavily towards 1-bp mutations, especially for insertions, as we did not observe a single insertion above 3 bp. We found 22 deletions greater than 3 bp in length, with the largest being 15 bp in length. None of the deletions greater than 1 bp in length were associated with homopolymers.

**Figure 5.**
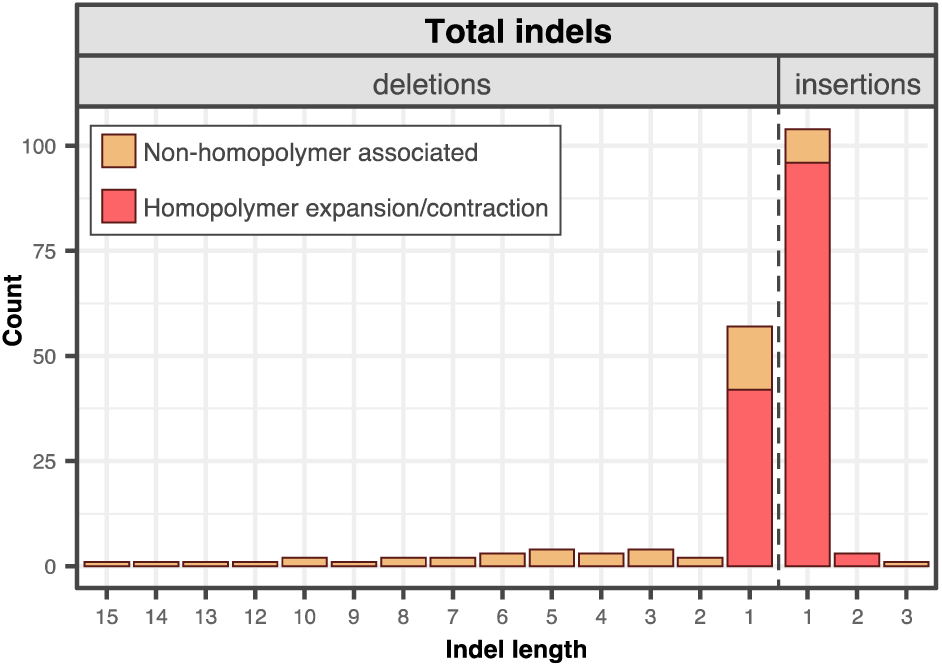
Indels are predominately found at homopolymers in *C. elegans* mtDNA. Here, indels that were shared between replicates were treated as independent events. See Figure S5 for counts that assume indels shared between replicates are the result of common ancestry. Indels that are expansions or contractions of existing homopolymers are shown in red, while those that are not associated with homopolymers are shown in tan.

## CONCLUSION

Duplex Sequencing is a promising alternative approach to labor and time intensive MA experiments for understanding mechanisms of mutation in metazoan mtDNAs. The idea that these genomes experience increased mutational loads given the proximity of the mtDNA to electron transport and associated ROS (Harman 1972) predates initial observations that mtDNA mutation rates are elevated above nuclear mutation rates in metazoans (Brown *et al*. 1979). However, numerous experimental studies have not supported the hypothesis that oxidative damage drives rapid mtDNA evolution (Halsne *et al*. 2012; Kennedy *et al*. 2013; Itsara *et al*. 2014). The large number of variants we detected with Duplex Sequencing provide a high-resolution view of the *C. elegans* mtDNA mutation spectrum (Fig 1a). These data support a role for oxidative damage because one of the predominant mutation classes, CG→AT transversions, is considered to be a hallmark of oxidative damage (Kennedy *et al*. 2013; Kino *et al*. 2017; Krašovec *et al*. 2017; Poetsch *et al*. 2018). In addition to CG→AT transversions, the spectrum also contains a high frequency of CG→TA transitions, and both classes show significant strand asymmetries (Fig 1b), with C→T and G→T changes enriched on the F-strand, providing further support that *C. elegans* mtDNA mutations are driven by single-stranded DNA damage. In contrast, if CG→TA transitions and CG→AT transversions were prevalent as a consequence of Pol γ misincorporations, it is not clear how this would lead to the strand asymmetries observed here. Further investigation is warranted to understand why CG→AT transversions are so abundant in the mtDNA of *C. elegans*, but relatively deplete in other metazoan mtDNAs. A surprising finding from recent experiments in mice (Kauppila *et al*. 2018) and fruit flies is that CG→AT transversions remain rare even in animals with deficiencies in BER (the principal pathway for repair of damaged DNA). Considering the high rate of CG→AT transversions observed here, it would be interesting to repeat this study with *C. elegans* lines lacking BER capabilities.

## DATA AVAILABILITY

The raw reads are available via the NCBI Sequence Read Archive (SRA) under accessions SRR14352240-14352248 (Duplex Sequencing libraries; Table S1) and SRR14352249, SRR14352237, and SRR14352238 (shotgun libraries 1, 2 and 3, respectively). The code used to process the raw reads, create consensus sequences and call variants is available here: https://github.com/dbsloan/duplexseq. The variants we detected through Duplex Sequencing are reported in File S1 (SNVs) and File S2 (indels).

## ACKNOWLEDGEMENTS

We thank Tai Montgomery for supplying N2 nematodes, media and lab space. We also thank Anne Hess for help in implementing the mixed linear models used in this study. This work was supported by the National Institutes of Health (R01 GM118046) and a National Science Foundation graduate fellowship (DGE-1450032).

## SUPPLEMENTARY MATERIAL

**Figure S1.**
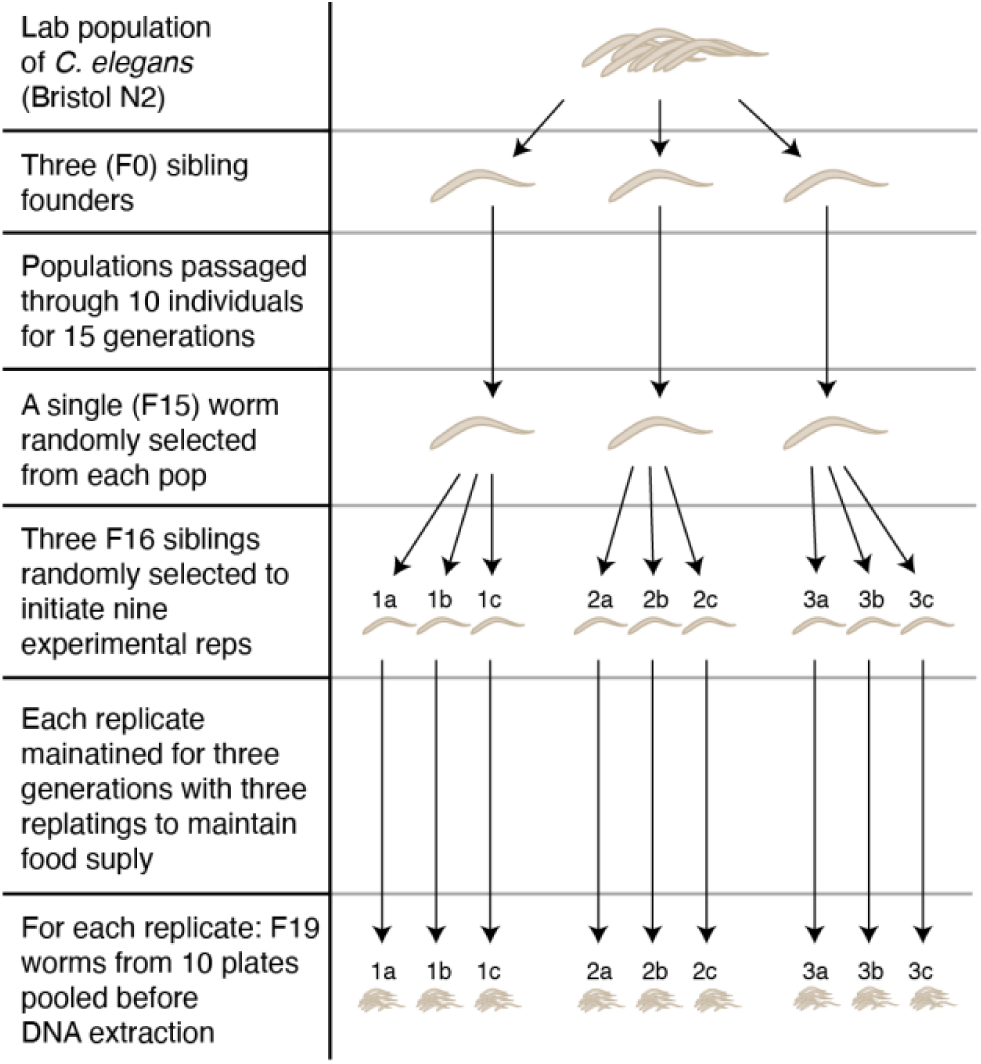
Culturing design used to obtain the nine replicates assayed in this study.

**Figure S2.**
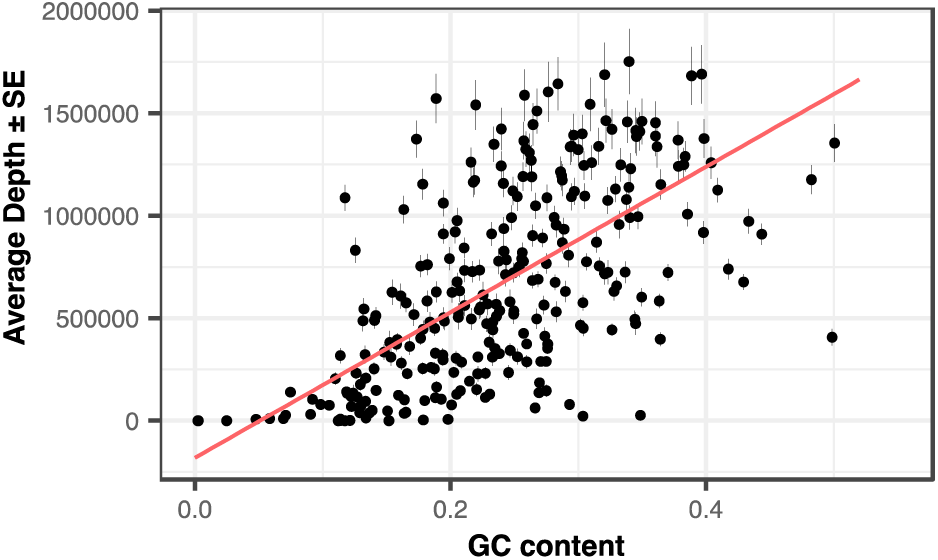
Average DCS depth as a function of GC content. Each point on the plot depicts a 50-bp window, with DCS depths averaged across the nine replicates and error bars reporting one standard error. Points were jittered for clarity.

**Figure S3.**
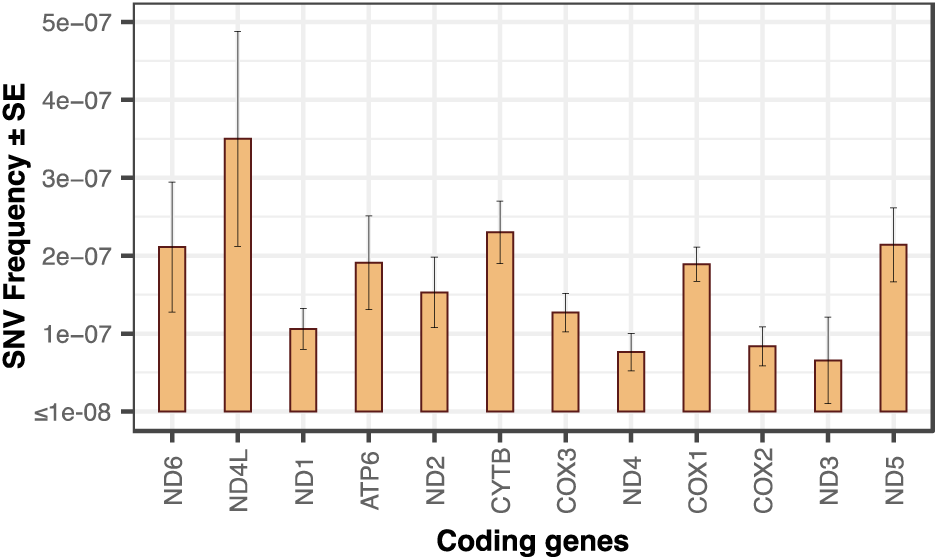
Variation in total SNV frequencies across the 12 protein-coding genes for all six substitution classes (one way ANOVA, *p* = 0.0072). In separate tests with each substitution class, significant between gene variation was observed only for only CG→AT transversions and CG→TA transitions (see Figure 2c).

**Figure S4.**
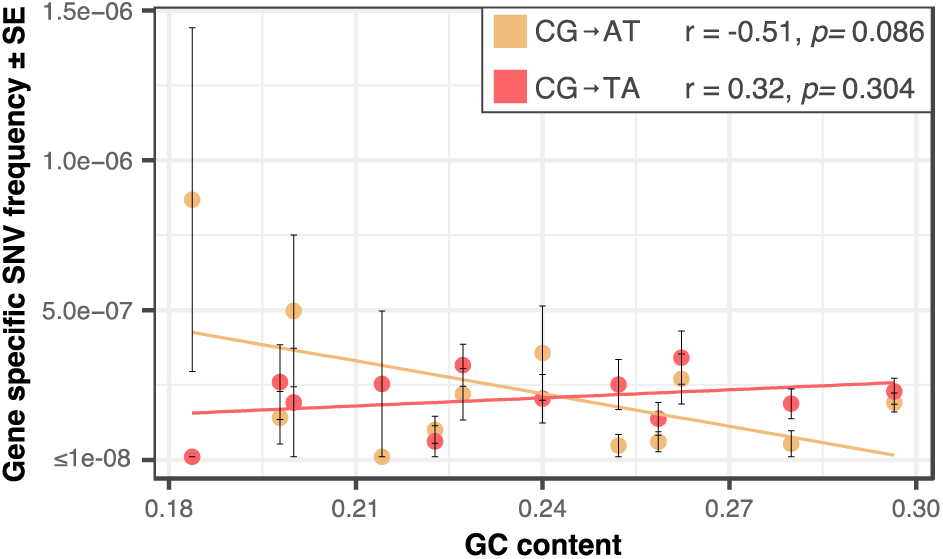
Correlation between gene specific SNV frequencies and GC content for CG AT transversions and CG TA transitions. R and *p* values (shown in legend) are from a Pearson correlation, implemented in R with the cor.test command.

**Figure S5.**
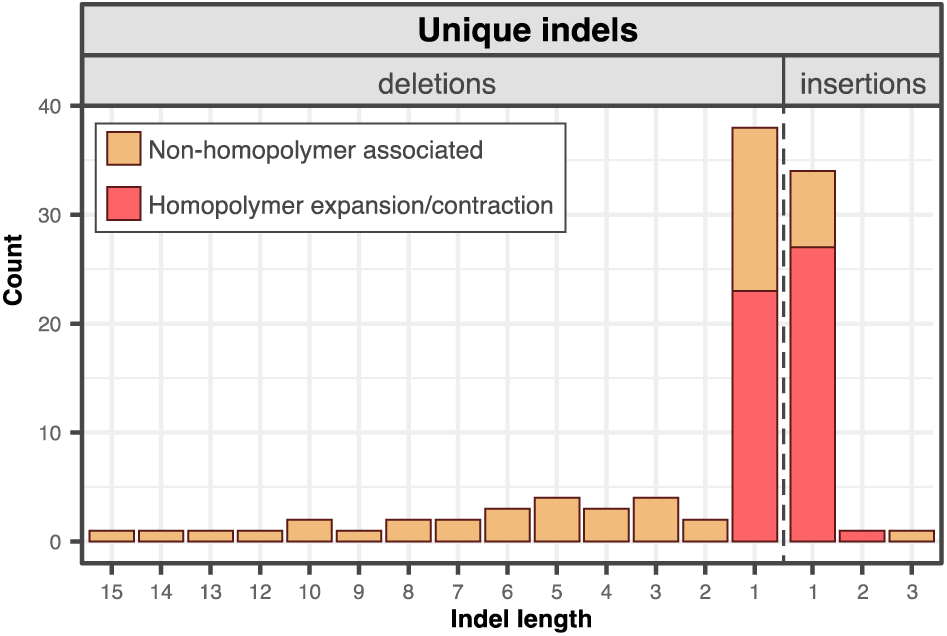
As in Figure 5, except that shared indels are assumed to be shared due to common ancestry, so are only counted once regardless of the number of replicates in which they occur.

**Table S1.**
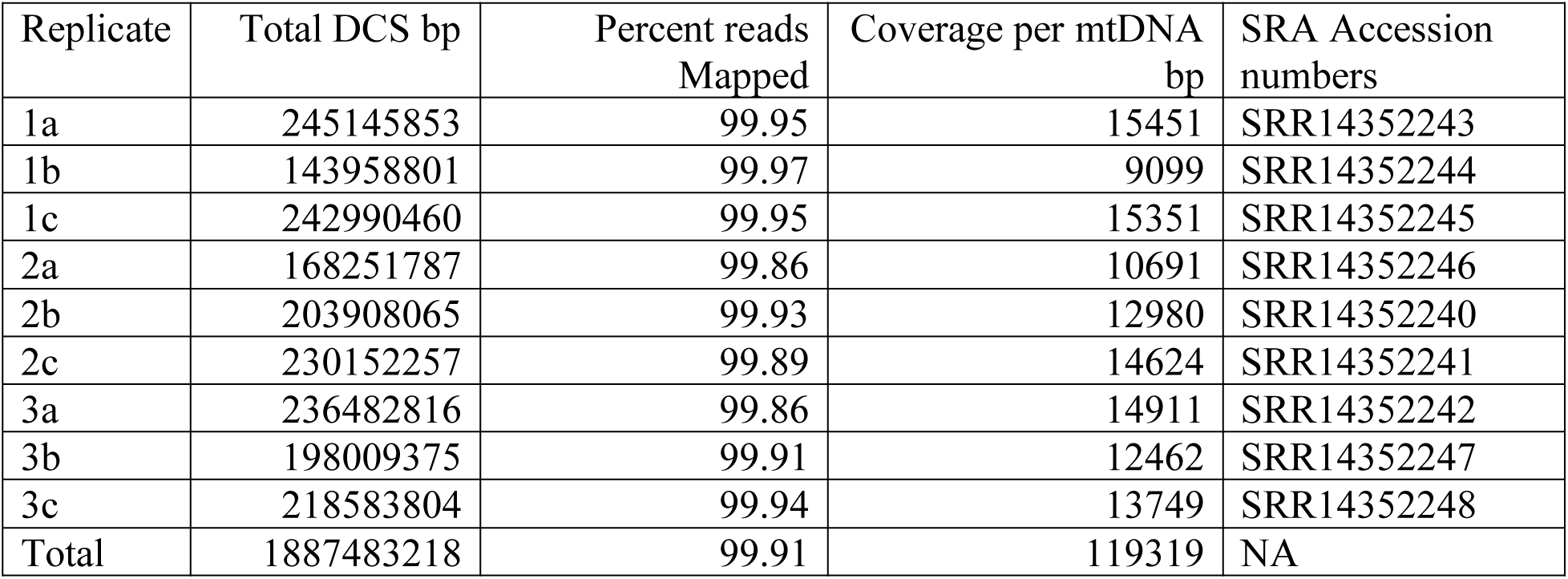
DCS coverage and percent mapping for each replicate.

| Replicate | Total DCS bp | Percent reads Mapped | Coverage per mtDNA bp | SRA Accession numbers |
| --- | --- | --- | --- | --- |
| 1a | 245145853 | 99.95 | 15451 | SRR14352243 |
| 1b | 143958801 | 99.97 | 9099 | SRR14352244 |
| 1c | 242990460 | 99.95 | 15351 | SRR14352245 |
| 2a | 168251787 | 99.86 | 10691 | SRR14352246 |
| 2b | 203908065 | 99.93 | 12980 | SRR14352240 |
| 2c | 230152257 | 99.89 | 14624 | SRR14352241 |
| 3a | 236482816 | 99.86 | 14911 | SRR14352242 |
| 3b | 198009375 | 99.91 | 12462 | SRR14352247 |
| 3c | 218583804 | 99.94 | 13749 | SRR14352248 |
| Total | 1887483218 | 99.91 | 119319 | NA |

**Table S2.** Positions which differ between the published N2 mitochondrial genome (NC_001328.1) and our lab N2 line. The two SNPs were completely fixed compared to NC_001328.1. In contrast, the 10 bp indel was supported by the majority of all DCSs, but it was not completely fixed in any of the nine replicates.

| Type of difference | Position in NC_001328.1 | Reference base in NC_001328.1 | Reference base in our lab N2 line |
| --- | --- | --- | --- |
| SNP | 8429 | A | G |
| SNP | 12998 | C | T |
| indel | 3235 | A<sub>11</sub> (11 bp homopolymer) | A<sub>10</sub> (10 bp homopolymer) |

**Table S3.**
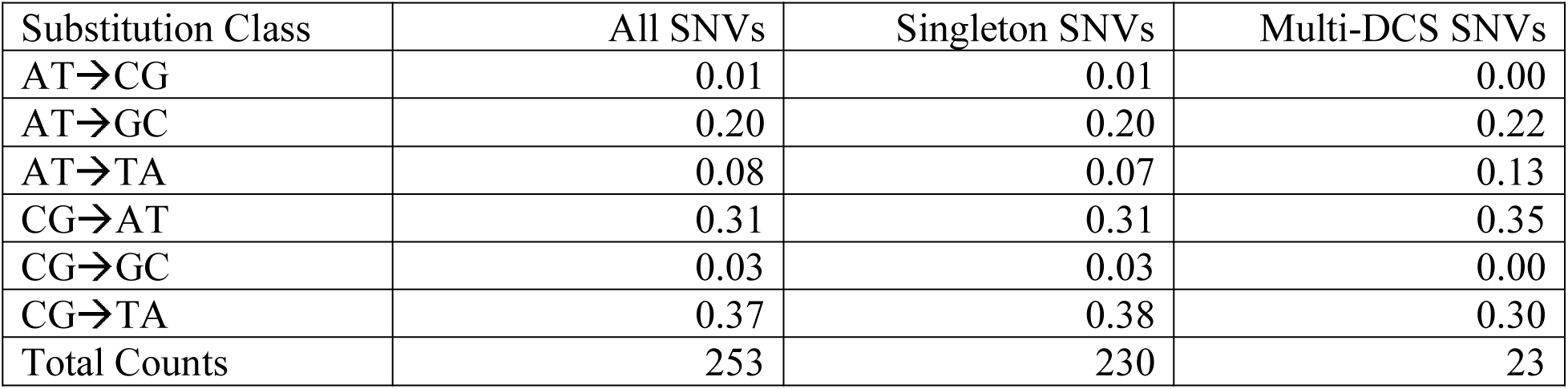
Proportions of the counts of each substitution class of the total counts for all SNVs, singleton SNVs, and multi-DCS SNVs. Note these proportions are not normalized for biased base composition of the *C. elegans* mtDNA or for differential probability of detection for AT and CG increasing variants due to uneven AT and CG coverage. For a normalized spectrum, see Figure 1.

**Table S4.** Proportions of the counts of each substitution class of the total counts for all SNVs as well as from the comparison of mtDNAs from 38 C. elegans natural isolates (Thompson *et al*. 2013; Konrad *et al*. 2017). Because substitutions in the population data cannot be reliably polarized, all substitution classes have been collapsed into four reversable classes (one transition and three transversions). As in Table S2, the proportions are not normalized to reflect the base composition of the mtDNA.

| Substitution Class | All SNVs | Natural isolates all substitutions | Natural isolates substitutions at four-fold degenerate sites |
| --- | --- | --- | --- |
| AT↔GC | 0.57 | 0.83 | 0.77 |
| AT↔CG | 0.32 | 0.07 | 0.09 |
| AT↔TA | 0.08 | 0.10 | 0.15 |
| GC↔CG | 0.03 | 0.002 | 0.00 |
| Total Counts | 253 | 408 | 162 |

